# Soil fauna may buffer the negative effects of drought on alien plant invasion

**DOI:** 10.1101/2021.07.26.453896

**Authors:** Huifei Jin, Liang Chang, Mark van Kleunen, Yanjie Liu

## Abstract

Although many studies have tested the direct effects of drought on alien plant invasion, less is known about how drought affects alien plant invasion indirectly via other groups of organisms such as soil fauna. To test for such indirect effects, we grew single plant of nine naturalized alien target species in pot-mesocosms with a native community of five native grassland species under four combinations of two drought (well-watered *vs* drought) and two soil-fauna-inoculation (with *vs* without) treatments. We found that drought decreased the absolute and the relative biomass production of the alien plants, and thus reduced their competitive advantage in the native community. Drought decreased the abundance of soil fauna, particularly the soil mites, but did not affect abundance and richness of soil herbivores. Soil-fauna inoculation increased the biomass of the native plant community and thereby decreased the relative biomass production of the alien species. The increased invasion resistance due to soil fauna, however, tended (*p* = 0.09) to be stronger for plants growing under well-watered conditions than under drought. Our multispecies experiment thus shows that soil fauna might help native resident communities to resist alien plant invasions, but that this effect might be diminished by drought.

## INTRODUCTION

With the rapid globalization, more and more plant species have been introduced to new regions outside their native range [1–3]. Some of these alien plants have become invasive, which could decrease native species diversity, change nutrient cycles, and thereby affect ecosystem functions [4–8]. Forecasts show that the number of alien plant species per continent may increase on average by 18% from 2005 to 2050 [9], indicating that the impacts of plant invasions on ecosystems may become even more severe. However, as there are many potential drivers of invasions, there are many uncertainties about how invasions and their impacts will develop in the future [10]. In this regard, particularly the potential effects of ongoing climate change on plant invasions have garnered interest [11, 12].

In many places, climate change is characterized by more frequent and more intense drought events [13–16]. Droughts do not only reduce water availability to the plants, but also decreases the nutrient absorption [17, 18]. Consequently, drought might affect the competition between alien and native plants [17, 19, 20], and thereby affect invasion success of alien plants [17, 21, 22]. For example, Manea et al. found that drought could reduce biomass production of native grasses, and consequently enhance the establishment success of alien plants [19]. However, many other studies also found that invasive plants may suffer more from drought than native species, indicating that drought could also suppress alien plant invasion [12, 20, 23–26]. One of the reasons for the mixed findings could be that most studies only considered direct effects of drought on alien plant invasion [12, 24, 26, 27]. Much less is known about the indirect effects, i.e. how drought affects alien plant invasion via other trophic levels.

Soil fauna include several important belowground trophic levels, which are increasingly recognized to affect plant competition [28–31]. Some of them could enhance nutrient mineralization, and thereby increase plant nutrient uptake [28, 32–37]. Given that invasive plants frequently respond more positively to nutrient enrichment than native plants [12, 38], the higher nutrient availability caused by soil fauna may increase the invasion success. In addition, soil fauna can also change alien-native competition via herbivory effects [39, 40]. The enemy-release hypothesis poses that alien plants are released from most of their native enemies [41–44]. Following this logic, alien plants would be damaged less than natives by herbivorous soil fauna, and therefore soil fauna would promote alien plant invasion. However, until now very few studies have tested how soil fauna affects alien plant invasion into resident communities [39, 40].

It has been found that indirect effects of altered biotic interactions due to climate change on animal populations are more pronounced than their direct effects [45], which may also be the case for plant populations. For example, empirical studies indicated that belowground trophic interactions could alter plant responses to drought [46–49]. Consequently, it is likely that drought might indirectly affect alien plant invasion in resident communities via effects on soil fauna. Indeed, drought can reduce the abundance and diversity of soil fauna [47, 50–53]. Given that soil fauna, in particular some of the root herbivores might affect competition between native and alien plants, indirect effects of drought on alien plant invasion in resident community would occur via the cascading effect of drought suppression on soil fauna [39, 40]. However, it has not been tested yet how drought could indirectly, via effects on soil fauna, affect plant invasion.

To test the direct and indirect effects (i.e. via soil fauna) of drought on alien plant invasion into a native resident community, we performed a mesocosm-pot experiment. We grew single plants of nine alien target species in a community of five native grassland species under four combinations of two drought (well-watered *vs* drought) and two soil-fauna-inoculation (with *vs* without) treatments. By comparing the absolute aboveground biomass production of the alien target species as well as their biomass production relative to the biomass production of the native competitors, we addressed the following specific questions: (1) Does drought suppress the absolute and relative biomass of alien species? (2) Does the presence of soil fauna promote or suppress the absolute and relative biomass of alien species? (3) Does the presence of soil fauna change the effect of drought on the absolute and relative biomass of alien species?

## MATERIAL AND METHODS

### Study species

To test the effects of drought, the presence of soil fauna, and their interaction on alien plant invasion in a native grassland community, we chose nine naturalized alien species as targets and five native species as competitors from the herbaceous flora of China (see Table S1). We classified the species as naturalized alien or native to China based on information in the book “The Checklist of The Naturalized Plants in China” [54] and the Flora of China database (https://www.efloras.org). To cover a wide taxonomic breadth, the nine alien species were chosen from eight genera of four families. The five native species, used to create the native community, included two forbs and three grasses that are all very common and do co-occur in many grasslands of China. Seeds of all species, except one, whose seeds were bought from a commercial seed company, were collected from natural populations growing in grasslands (Table S1).

### Soil-fauna collection

To provide a live soil-fauna community as inoculum for the pot mesocosms, we collected soil fauna from a grassland site, where the five native species used for present study also occur in, at the Northeast Institute of Geography and Agricultural Ecology, Chinese Academy of Sciences (125°24′30″E, 43°59′49″N) on 21 July 2020. In the grassland, we removed aboveground plant materials from each of 100 sampling locations (30cm × 30cm), and then collected from each location a soil sample of 1-L (10cm × 10cm× 10cm) using a shovel. Each sampling location was at least 10m apart from the others. Then, we brought the 100 soil samples back to the laboratory, where we extracted the soil fauna communities of each soil sample separately using Berlese-Tullgren extractors without heating (https://en.wikipedia.org/wiki/Tullgren_funnel). In brief, we put all soil samples on top of 100 stainless steel soil-sieves with a 2mm mesh size, and then waited 12 days so that many of the soil organisms would fall through the holes, via stainless steel funnels, into plastic bottles filled with 50 ml soil-fauna-free peat moss (Pindstrup Plus, Pindstrup Mosebrug A/S, Denmark). On 2 August 2020, we finished the soil-fauna-community collection. We then randomly chose ten of the 100 soil-fauna communities for soil-fauna investigation (see Table S2), and used the remaining 90 soil-fauna communities for inoculating of the soil in the experimental pot mesocosms.

### Experimental set-up

To compare the growth performance of alien plants when growing in a resident native grassland community under different drought and soil-fauna-inoculation treatments, we did a mesocosm-pot experiment in a greenhouse of the Northeast Institute of Geography and Agricultural Ecology, Chinese Academy of Sciences. We grew each of the nine alien species in the centre of a matrix of the native community under two water availabilities (well-watered *vs* drought) and two soil-fauna-inoculation (with *vs* without) treatments.

From 15 May to 5 July 2020, we sowed the seeds of each species separately into plastic trays (195 mm × 146 mm × 65 mm) filled with peat moss as substrate (Pindstrup Plus, Pindstrup Mosebrug A/S, Denmark). As previous experiments had shown that the time required for germination differs among the species, we sowed the species on different dates (Table S1) to obtain similarly sized seedlings at the start of the experiment. On 3 August 2020, we filled 180 2.5-L circular plastic pots (top diameter × bottom diameter × height: 18.5 × 12.5 × 15 cm, Yancheng Tengle Plastics Co., Ltd, China) with the same substrate as used for germination. To avoid nutrient limitation during plant growth, we mixed each pot with 5 g slow-release fertilizer (Osmocote® Exact Standard, Everris International B.V., Geldermalsen, The Netherlands). To create the native community, we selected similarly sized seedlings from each of the five native species, and transplanted one seedling of each native species at equal distances in a circle (diameter = 11 cm) around the centre of each pot. We then planted into the centre of each pot one seedling of one of the alien species. For each of the nine alien species, we had 20 pots.

After transplanting, we randomly assigned two pots of each alien species to each of ten plastic cages (150 cm × 90 cm × 100 cm; Figure 1). The ten cages were covered with nylon nets (mesh size: 0.15 mm × 0.15 mm) to keep out small soil fauna. We put a plastic dish under each pot, and regularly watered the pots before starting the drought treatment to ensure that none of the plants were water limited. On 3 August 2020, we inoculated each pot in five of the ten cages (i.e. 90 pots) with 50 ml of the soil-fauna inoculum we had collected. As the soil-fauna inoculum also introduced another 50 ml of peat moss into pots, we also added 50 ml of the same peat moss (free of soil fauna) to each pot of the remaining five cages as a control. We assigned the cages with and without soil-fauna inoculations to alternating positions that were at least 0.5 m apart from each other (Figure 1). On 30 August 2020 (i.e. 27 days after the start of the experiment), we started the drought treatment. One of the two pots of each alien target species in each cage served as a control, which was watered regularly to keep the substrate moist throughout the entire experiment, while the other pots did not receive water unless the plants were wilted. We daily checked all pots of the drought treatment, and when some plants of the community in a pot were wilted (i.e. had lost leaf turgor), we supplied the pot with 50 ml of water. So, for each of the nine alien species, we had five replicates of each of the four drought × soil-fauna-inoculation treatment combinations, resulting in a total of 180 pots (9 alien species × 2 soil-fauna inoculations [with *vs* without] × 2 drought treatments [well-watered *vs* drought] × 5 replicates [i.e. cages]).

**Figure 1.**
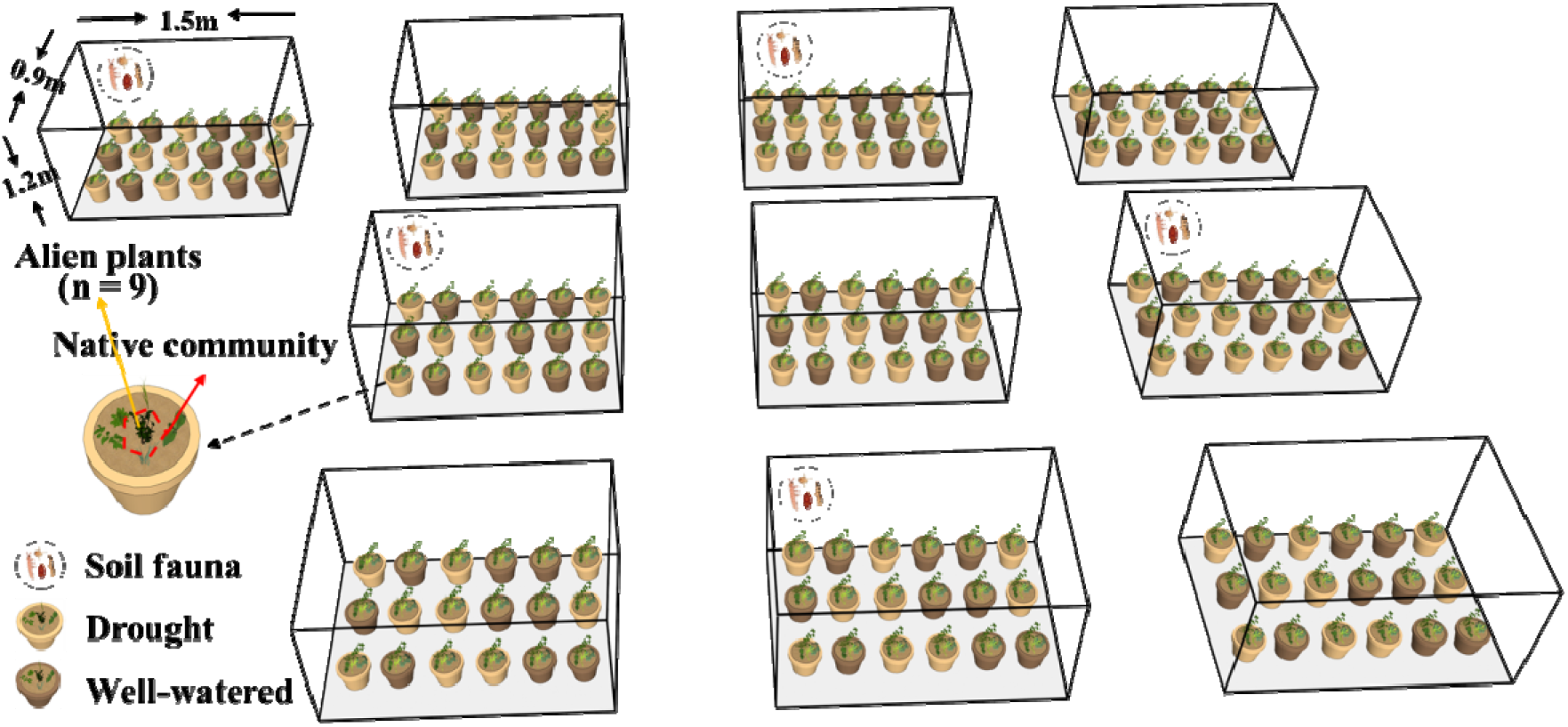
Graphical illustration of the experimental design.

On 28 September 2020, 10 weeks after the start of the drought treatments, we harvested the experiment. For each pot, we separately harvested the aboveground biomass of the alien target species and of the native community. As the roots of the plants were too much intertwined, we could not harvest the belowground biomass. All aboveground biomass was dried for at least 72 h at 65 °C, and then weighed. Based on the aboveground biomass of the alien and native species, we calculated the biomass proportion of the alien target species (the biomass of the alien target species / [biomass of the alien target species + biomass of the native community]) as a proxy of the dominance of the alien target species (for a similar approach, see Refs. 38, 55).

To test the effect of the drought treatment on soil fauna, we investigated the soil fauna of drought and well-watered pots after plant harvest. We first randomly selected one pot of each target alien species under drought or well-watered conditions, and then brought all soil of these pots back to the laboratory. In total, we had 18 soil samples for soil-fauna extraction (9 alien species × 2 drought treatments [well-watered *vs* drought] × 1 replicate). Using the Berlese-Tullgren extractors without heating, we extracted the soil fauna communities of each soil sample separately, and obtained of the total soil-fauna abundance, soil-mites abundance, soil-herbivore abundance and soil-herbivore richness for each sample. As most soil-fauna species were mites (*c*. 70.0%) for our samples, and it was difficult to identify all of them to species level, we do not have data of soil-fauna richness. We also did soil-fauna investigation for five pots that were randomly selected from the pots without soil-fauna inoculation, and found no soil fauna in these pots, indicating that our soil-fauna inoculation treatment and the isolation imposed by the cages were effective.

### Statistical analysis

To analyze the effects of drought, soil-fauna-inoculation treatments and their interactions on performance of the alien plants in the native community, we fitted linear mixed-effects models using the lme function of the ‘nlme’ package [56] in R 4.0.3 [57]. Aboveground biomass production of the alien target species, the native competitor species and biomass proportion of the alien target species in each pot (i.e. target aboveground biomass/total aboveground biomass) were the response variables. To meet the assumption of normality, biomass production of the alien target species and the native competitor species were natural-log-transformed, and biomass proportion of the target species was logit-transformed. We included drought treatment (i.e. well-watered *vs* drought), soil-fauna-inoculation treatment (i.e. with *vs* without addition) and their interactions as fixed effects in all models.

To account for non-independence of individuals of the same alien plant species and for phylogenetic non-independence of the species, we included identity of the target species nested within family as random factors in all models. In addition to account for non-independence of plants within the same cage, we also included cage identity as a random factor in all model. As the homoscedasticity assumption was violated in all models, we also included variance structures to model different variances per species or per cage (based on model selection) using the “*varIdent*” function in the R package ‘nlme’ [56]. We used log-likelihood ratio tests to assess significance of the fixed effects drought treatment, soil-fauna-inoculation treatment and their interaction [58]. These tests were based on comparisons of maximum-likelihood models with and without the terms of interest, and the variance components were estimated using the restricted maximum-likelihood method of the full model [58]. To test the effect of drought on total soil-fauna abundance, soil-mites abundance, soil-herbivore abundance and soil-herbivore richness, we applied Kruskal-Wallis test using the kruskal.test function of the ‘stats’ package in R 4.0.3 [57].

## RESULTS

The specific response patterns of each alien species to the drought and soil-fauna-inoculation treatments were more or less similar with the general patterns of alien target species (Figure 2 & S1). Averaged across the nine alien target species, drought significantly decreased the aboveground biomass production of alien target species (−58.6%; Table 1; Figure 2a), and the native community (−51.5%; Table 1, Figure 2b). As the biomass of the aliens decreased more strongly in response to drought than the biomass of the natives did, the biomass proportion of the alien target species in each pot decreased (−11.6%; Table 1; Figure 2c). Inoculation with soil fauna had no significant effect on aboveground biomass of the alien target plants, but had a significant positive effect on aboveground biomass of the native community (+40.1%; Table 1; Figure 2b). Consequently, the aboveground biomass proportion of the alien target species was decrease in the presence of soil fauna (−41.9%; Table 1; Figure 2c). Moreover, as the native community benefitted more from the presence of soil fauna under well-watered conditions (+44.0%) than under drought conditions (+32.5%; Table 1; Figure 2b), the negative effect of soil-fauna inoculation on biomass proportion of the alien target species tended to be stronger under well-watered conditions (−49.7%) than under drought conditions (−32.0%; marginally significant S × D interaction in Table 1, *p* = 0.09; Figure 2c). In addition, we found that drought significantly decreased the total abundance of soil fauna (−57.4%; Figure 3a) and soil mites (−54.6%; Figure 3b), but did not affect soil-herbivore abundance and richness (Figure 3c & 3d).

**Figure 2.**
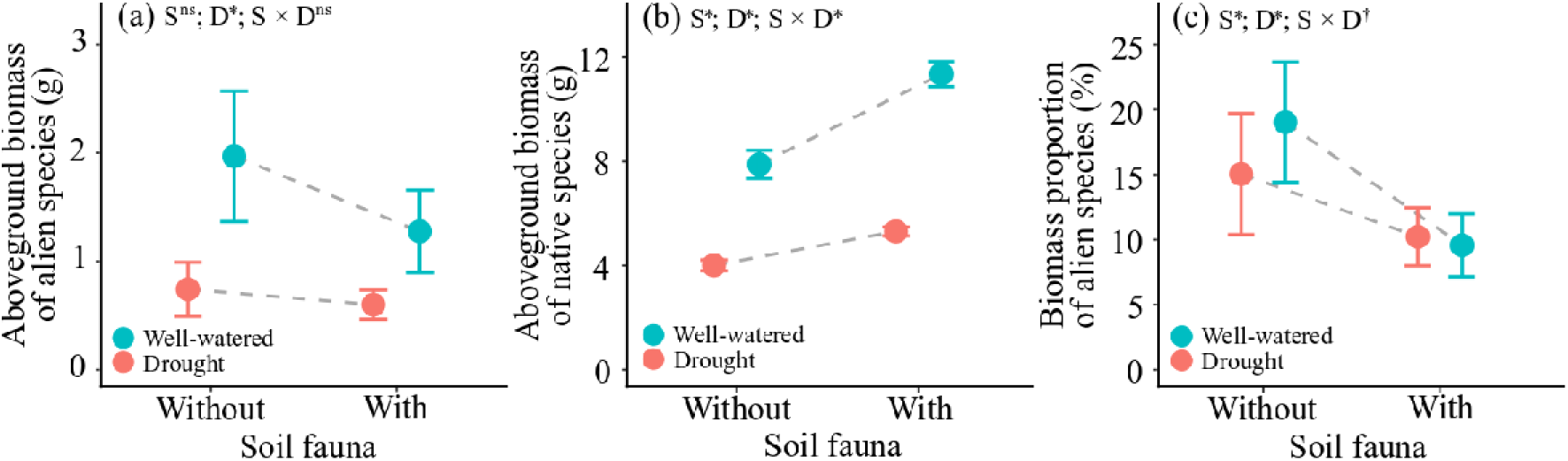
Mean values (± SE) of aboveground biomass of the alien target species (a), aboveground biomass production of the native competitor species (b), and biomass proportion of the alien target species in each pot (c) under different drought (well-watered *vs* drought) and soil-fauna-inoculation (with *vs* without) treatments. Significant parameters (i.e. ‘S’ [soil fauna], ‘D’ [drought] and ‘S × D’) are indicted with asterisks (*), the marginal significant ones are indicated with daggers (†), and the non-significant ones are indicated with “ns”.

**Table 1.** Results of linear mixed-effects models testing the effects of drought (well-watered *vs* drought), soil fauna (with *vs* without) and their interactions on aboveground biomass production of the alien target plants, native competitor species, and biomass proportion of the alien target species in each pot. Significant effects (*p* < 0.05) are in bold, and marginal significant effects (0.05 < *p* < 0.1) are underlined.

| Fixed effects | Aboveground biomass production of the<br>alien target species<br>(natural-log-transformed) |  |  | Aboveground biomass production<br>of the native competitor species<br>(natural-log-transformed) |  |  | Aboveground biomass proportion of the<br>target species in each pot<br>(logit-transformed) |  |  |
| --- | --- | --- | --- | --- | --- | --- | --- | --- | --- |
| | Df | $\chi^2$ | $p$ | Df | $\chi^2$ | $p$ | Df | $\chi^2$ | $p$ |
| Soil fauna (S) | 1 | 2.4990 | 0.1139 | 1 | 4.7722 | **0.0289** | 1 | 7.0803 | **0.0078** |
| Drought (D) | 1 | 63.8413 | **<0.0001** | 1 | 164.1043 | **<0.0001** | 1 | 5.6339 | **0.0176** |
| S × D | 1 | 0.1512 | 0.6974 | 1 | 5.6304 | **0.0177** | 1 | 2.8297 | <u>0.0925</u> |
| **Random effects** |  | ***SD*** |  |  | ***SD*** |  |  | ***SD*** |  |
| Family |  | 0.004 |  |  | 0.002 |  |  | 0.004 |  |
| Species |  | 0.970* |  |  | 0.086 |  |  | 1.057* |  |
| Cage |  | 0.004 |  |  | 0.267* |  |  | 0.187 |  |
| Residual |  | 0.586 |  |  | 0.126 |  |  | 0.555 |  |
|  |  | **Marginal $R^2$** | **Conditional $R^2$** |  | **Marginal $R^2$** | **Conditional $R^2$** |  | **Marginal $R^2$** | **Conditional $R^2$** |
| $R^2$ of the model | | 0.138 | 0.770 | | 0.641 | 0.940 | | 0.063 | 0.802 |

\* Standard deviations for individual alien species or individual cage random effects for the saturated model are found in Table S3.

**Figure 3.**
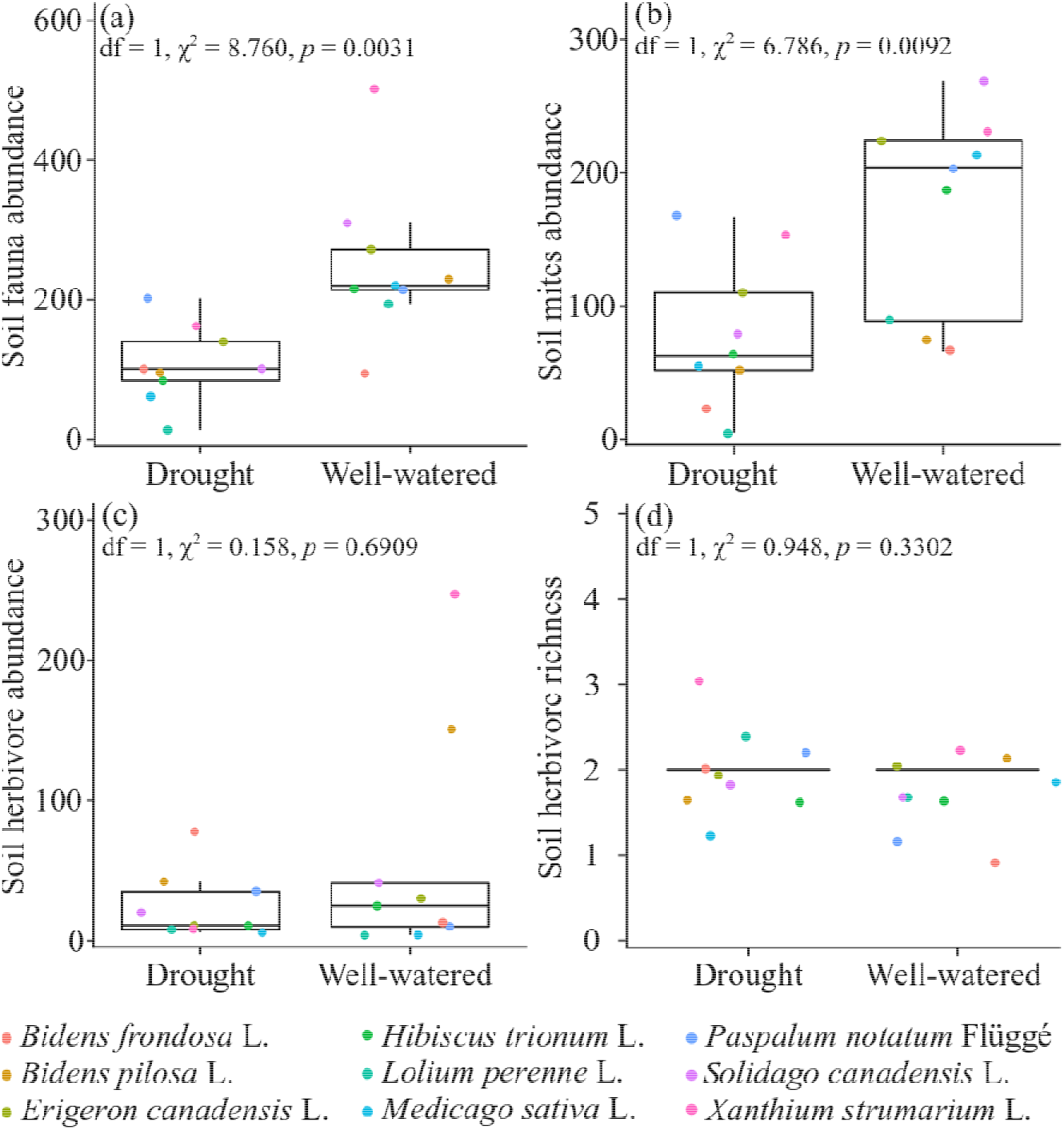
Total abundance of soil fauna (a), soil mites (b) and soil herbivores (c), as well as the soil herbivore richness (c) under drought and well-watered conditions. The nine different color points represent samples of the nine pots with different alien target species.

## DISCUSSION

Our multispecies experiment found that drought limited the absolute and the relative biomass production of the alien target plants. This means that drought suppressed the growth performance and thus reduced the competitive ability of the alien species in the native community. In addition, we found that the presence of soil-fauna communities benefited the native community and resulted in a decreased biomass proportion of the alien species. In other words, the presence of soil fauna promoted the resistance of the native community against invasion by the alien species. Moreover, the suppressive effect of drought on biomass proportion of the alien plants tended to disappear (although this effect was only marginally significant; *p* = 0.09) in the presence of soil fauna. This suggests that the presence of soil-fauna communities might negate the negative effect of drought on alien plant invasion into a resident community.

While drought is well known to reduce plant performance overall [59–61], recent studies found that growth and reproduction were more strongly affected for invasive than for native plant species [12, 23, 26]. Our results are consistent with these previous findings (see also the total biomass production per pot in Figure S2), and suggest that the native competitors were more tolerant to drought than the invasive alien species [12, 20, 23–26]. On the other hand, it could also indicate that the invasive alien plants took more advantage of the well-watered conditions than the native species did. This would be in line with the idea that invasive plants show higher phenotypic plasticity, and capitalize more strongly on benign conditions than native plants do (i.e. the Master-of-some strategy *sensu* Ref. 62). In any case, the negative effect of drought on the biomass proportion of the alien plants suggests that the competitive balance between invasive alien plants and native plants could be changed by drought in favour of the resident community. Another recent study, however, showed that alien species that are not invasive yet could benefit more from drought relative to resident plants [22], which could imply a turn-over in invasive alien species with ongoing climate change.

Inoculation with soil fauna significantly increased the biomass production of the native community. It is often suggested that soil fauna, such as collembolans and mites in our study (Table S2), could enhance soil nutrient mineralization and consequently nutrient absorption of plants [32, 63]. As a consequence, soil fauna frequently have positive effects on plant performance [63–65]. However, we found that soil-fauna inoculation had no statistically significant effect on growth of the alien target plants. Therefore, it is unlikely that soil fauna promoted native plant growth by increasing nutrient availability, as we would have expected the invasive alien species to benefit from it too. To confirm this, we further measured the total nitrogen and alkaline hydrolyzable nitrogen (i.e. *plant-available* nitrogen) of 36 soil samples (20% of total pots) from the drought treatment. This showed that soil-fauna inoculation did not significantly affect both types of soil nitrogen (Figure S3).

The most likely explanation for why soil-fauna inoculation did not affect alien plant growth is that the invasive alien species may have been released from their native enemies in the study area [66, 67]. In contrast, native plants were most likely attacked by the soil herbivores. The fact that they produced more aboveground biomass might indicate that the presence of soil fauna resulted in a re-allocation of biomass from belowground to aboveground structures or that compensatory growth actually resulted in overcompensation [68, 69]. Indeed, it has been previously shown that collembolan, at intermediate densities, could enhance plant growth through herbivory-induced overcompensation [70]. Although this could explain why the soil fauna suppressed the dominance (i.e. biomass proportion) of alien target plants, future explicitly experiments testing such soil-herbivory-induced overcompensation are needed. Nevertheless, while it has been shown before that soil fauna affects the composition of plant communities [28, 30, 39], this is one of the first studies to document the ability of the soil fauna to provide resistance against alien plant invasion.

Although numerous studies have shown that climate change could affect interactions of plants with aboveground organisms at other trophic levels [51, 71–73], only few studies have addressed how climate change may affect interactions of plants with belowground organisms at other trophic levels [47, 49, 51, 74]. To the best of our knowledge, no studies have addressed how climate change interacts with resident soil fauna to affect the competitions between native and invasive plant species. Many previous studies found that drought could decrease the abundance of soil fauna [50–52, 75, 76]. Our study supports this, and further showed that drought mainly decreased the abundance of soil mites rather than soil herbivores. Given that drought also decreased the growth performance of native plants, the constant soil-herbivore abundance would lead to a higher damage intensity, and thus the overcompensation decreased in response to drought. This could explain why the growth promotion of native plants induced by soil fauna was stronger under well-watered condition than under drought. Our present finding is also consistent with previous studies, showing that drought can hinder insect-induced-overcompensation of plants [68, 77].

Interestingly, we found tentative evidence that the decrease in dominance of alien target plant caused by drought was larger in the absence of soil fauna than in its presence. In other words, the presence of soil fauna might buffer against the negative effects of drought on alien plant invasion in resident communities. This is because soil fauna did not mediate the drought effects on the growth of alien plants which were most likely released from their native enemies. On the other hand, it amplified the negative effect of drought on the growth of native plants. Our study thus indicates that previous pot experiments that did not include soil fauna might have overestimated the effects of drought on alien plant invasion. Therefore, future studies testing effects of climate change on alien plant invasion should also consider the role of other belowground trophic levels.

## CONCLUSIONS

The findings of our multi-species experiment are in line with results of previous studies that drought can inhibit alien plant invasion into a native resident community. However, to the best of our knowledge, we here show for the first time that the presence of soil fauna might help the resident plant community to resist alien plant invasions. This soil-fauna mediated resistance may be partly negated by drought. This implies that with ongoing climate change, and more frequent droughts, invasive plants might be more likely to overcome the resistance provided by soil fauna.

## ACKNOWLEDGEMENTS

We thank Xue Zhang, Lichao Wang and Yanjun Li for their practical assistance. YL acknowledges funding from the Chinese Academy of Sciences (Y9B7041001).

## AUTHOR CONTRIBUTIONS

YL conceived the idea and designed the experiment. HJ and LC performed the experiment. HJ and YL analyzed the data. HJ and YL wrote the first draft of the manuscript, with further inputs from MvK and LC.

## DATA ACCESSIBILITY

Should the manuscript be accepted, the data supporting the results will be archived in Dryad and the data DOI will be included at the end of the article.

## SUPPORTING INFORMATION

**Table S1.**
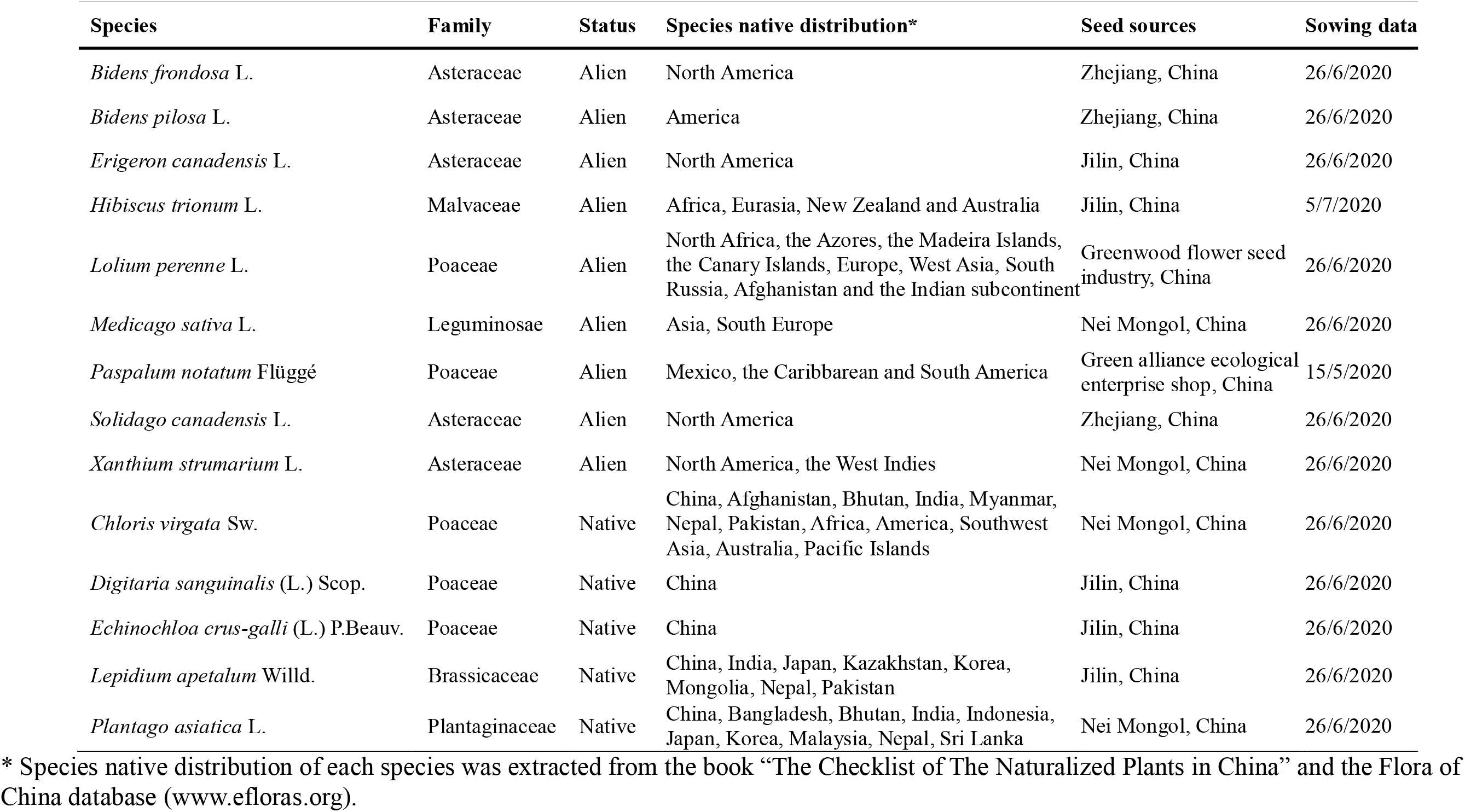
Details of the study species used in the experiment.

| Species | Family | Status | Species native distribution* | Seed sources | Sowing data |
| --- | --- | --- | --- | --- | --- |
| *Bidens frondosa* L. | Asteraceae | Alien | North America | Zhejiang, China | 26/6/2020 |
| *Bidens pilosa* L. | Asteraceae | Alien | America | Zhejiang, China | 26/6/2020 |
| *Erigeron canadensis* L. | Asteraceae | Alien | North America | Jilin, China | 26/6/2020 |
| *Hibiscus trionum* L. | Malvaceae | Alien | Africa, Eurasia, New Zealand and Australia | Jilin, China | 5/7/2020 |
| *Lolium perenne* L. | Poaceae | Alien | North Africa, the Azores, the Madeira Islands, the Canary Islands, Europe, West Asia, South Russia, Afghanistan and the Indian subcontinent | Greenwood flower seed industry, China | 26/6/2020 |
| *Medicago sativa* L. | Leguminosae | Alien | Asia, South Europe | Nei Mongol, China | 26/6/2020 |
| *Paspalum notatum* Flüggé | Poaceae | Alien | Mexico, the Caribbarean and South America | Green alliance ecological enterprise shop, China | 15/5/2020 |
| *Solidago canadensis* L. | Asteraceae | Alien | North America | Zhejiang, China | 26/6/2020 |
| *Xanthium strumarium* L. | Asteraceae | Alien | North America, the West Indies | Nei Mongol, China | 26/6/2020 |
| *Chloris virgata* Sw. | Poaceae | Native | China, Afghanistan, Bhutan, India, Myanmar, Nepal, Pakistan, Africa, America, Southwest Asia, Australia, Pacific Islands | Nei Mongol, China | 26/6/2020 |
| *Digitaria sanguinalis* (L.) Scop. | Poaceae | Native | China | Jilin, China | 26/6/2020 |
| *Echinochloa crus-galli* (L.) P.Beauv. | Poaceae | Native | China | Jilin, China | 26/6/2020 |
| *Lepidium apetalum* Willd. | Brassicaceae | Native | China, India, Japan, Kazakhstan, Korea, Mongolia, Nepal, Pakistan | Jilin, China | 26/6/2020 |
| *Plantago asiatica* L. | Plantaginaceae | Native | China, Bangladesh, Bhutan, India, Indonesia, Japan, Korea, Malaysia, Nepal, Sri Lanka | Nei Mongol, China | 26/6/2020 |

\* Species native distribution of each species was extracted from the book “The Checklist of The Naturalized Plants in China” and the Flora of China database (www.efloras.org).

**Table S2.**
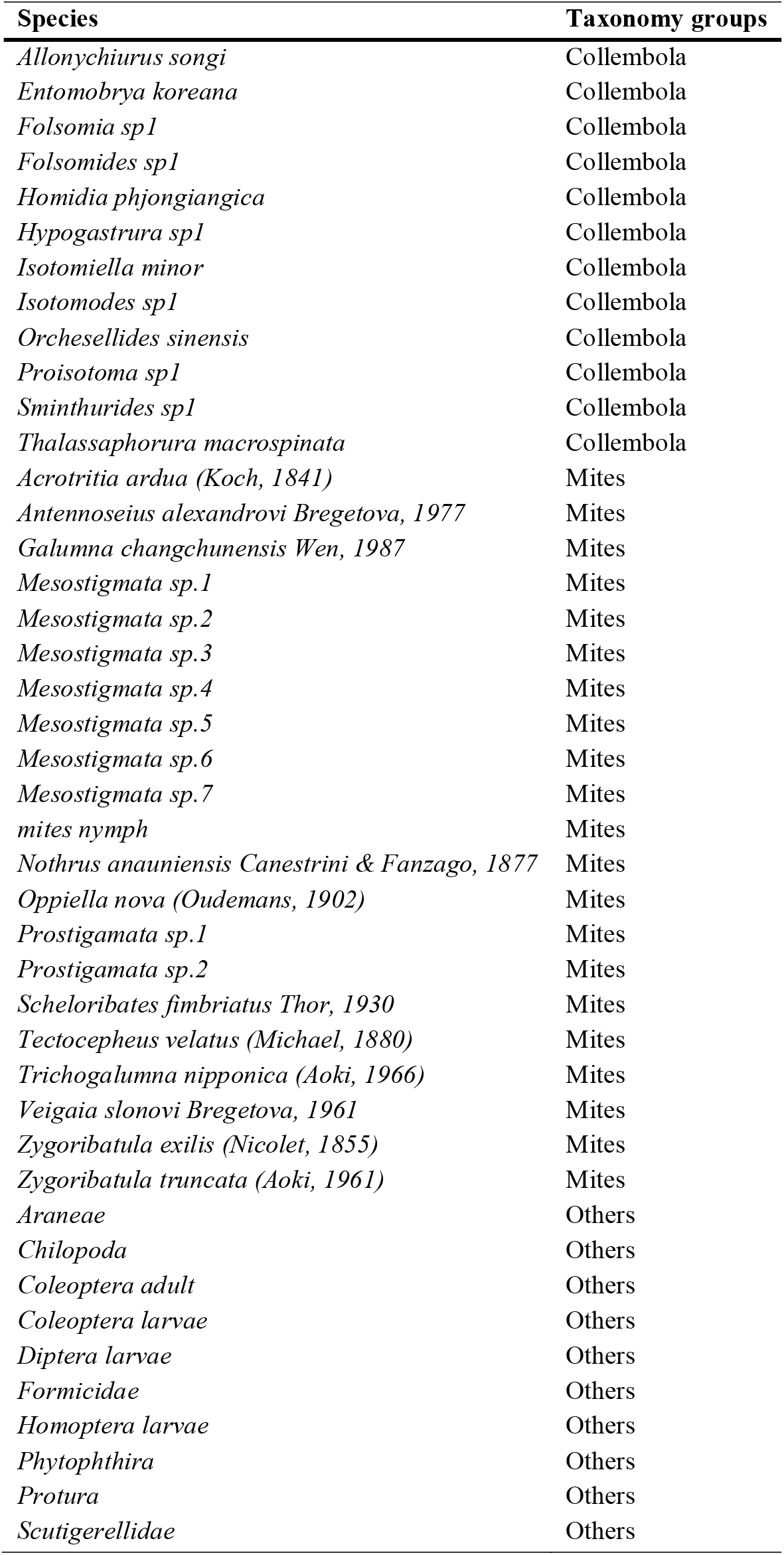
Soil-fauna taxa found in the soil used to inoculate the pot mesocosms.

**Table S3.** Standard deviations for individual alien species or individual cage random effects for metrics analyzed with models with a Gaussian error distribution. The standard deviations given refer to the first species and cage respectively. For each species and cage, these should be multiplied by the multiplication factors. The names of the alien species in the table are abbreviated using the first letter of the genus and species epithet.

| Metric | Species Standard Deviation | Multiplication factor standard deviation |  |  |  |  |  |  |  |  |
| --- | --- | --- | --- | --- | --- | --- | --- | --- | --- | --- |
|  |  | LP | EC | HT | PN | SC | BF | BP | MS | XS |
| Aboveground biomass production of the alien target species | 0.970 | 1.000 | 0.778 | 0.626 | 0.820 | 1.078 | 1.935 | 2.600 | 1.890 | 1.778 |
| Aboveground biomass proportion of the target species in each pot | 1.057 | 1.000 | 0.879 | 0.920 | 0.942 | 1.179 | 2.164 | 2.958 | 1.905 | 1.815 |

| Metric | Cages Standard<br>Deviation | Multiplication factor for standard deviation |  |  |  |  |  |  |  |  |  |
| --- | --- | --- | --- | --- | --- | --- | --- | --- | --- | --- | --- |
|  |  | Cage1 | Cage2 | Cage3 | Cage4 | Cage5 | Cage6 | Cage7 | Cage8 | Cage9 | Cage10 |
| Aboveground biomass production of the native competitor species | 0.267 | 1.000 | 2.461 | 2.603 | 3.751 | 1.531 | 1.725 | 2.090 | 1.302 | 3.503 | 2.590 |

**Figure S1.**
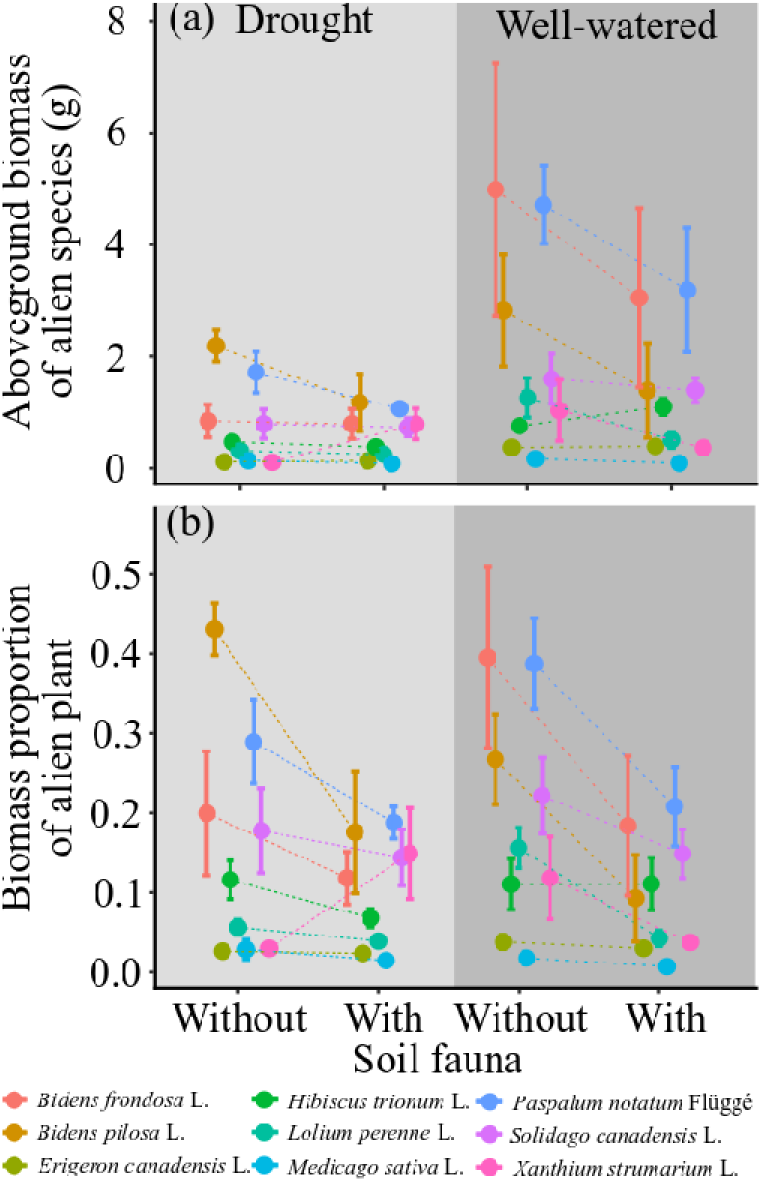
Mean values (± SE) of aboveground biomass of the specific alien target species (a) and biomass proportion of the specific alien target species in each pot (b) under different drought (well-watered vs drought) and soil-fauna-inoculation (with vs without) treatments.

**Figure S2.**
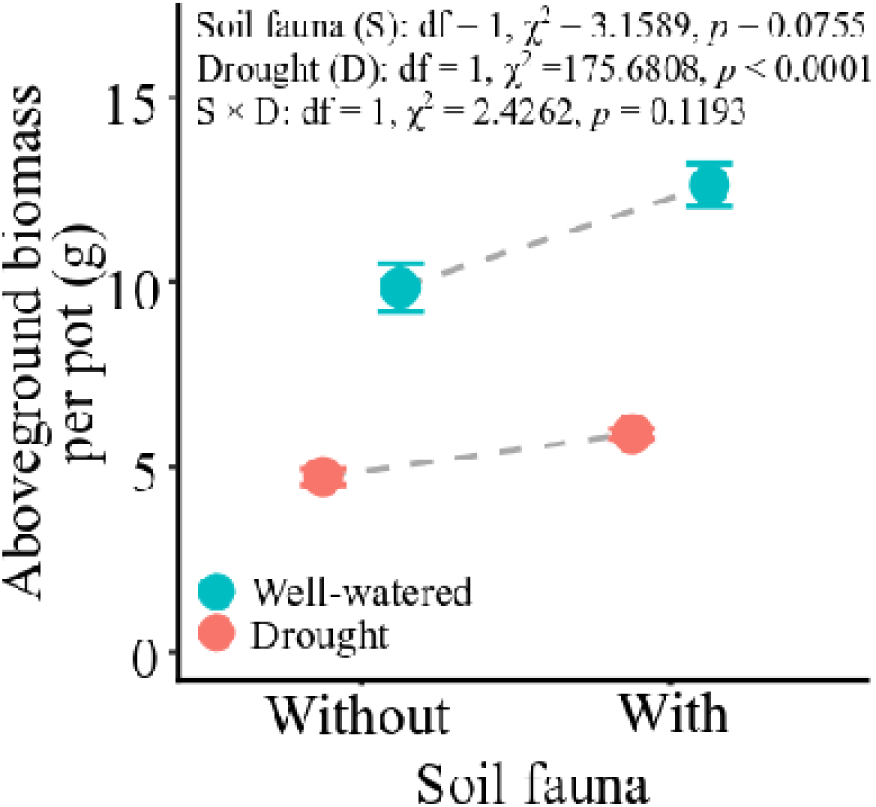
Mean values (± SE) of the total aboveground biomass production per pot (i.e. alien + native) under different drought (well-watered *vs* drought) and soil-fauna-inoculation (with *vs* without) treatments.

**Figure S3.**
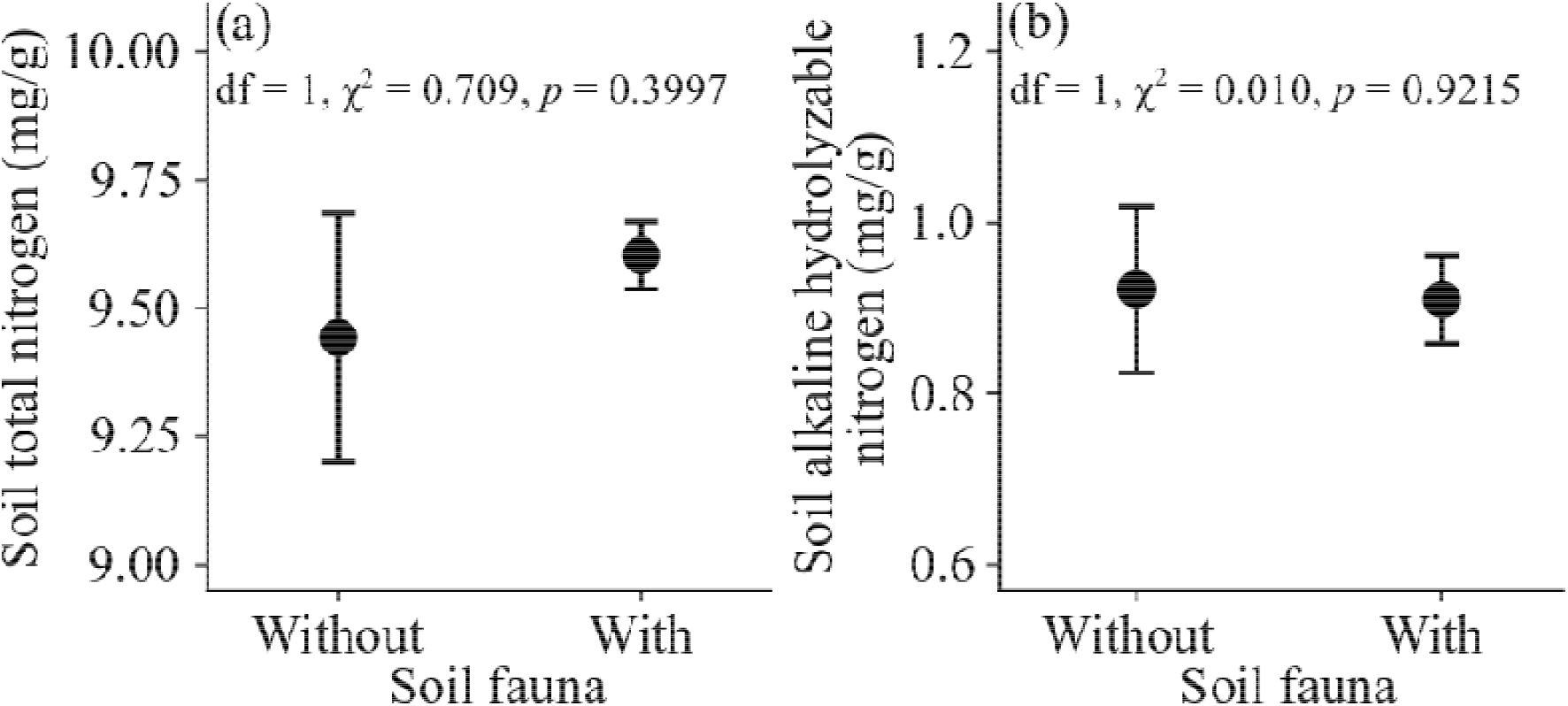
Mean values (± SE) of the soil total nitrogen (a) and soil alkaline hydrolyzable nitrogen (b) in the pots with and without soil-fauna-inoculation treatments.

## REFERENCES

[1] Seebens, H., Blackburn, T.M., Dyer, E.E., Genovesi, P., Hulmeh, P.E., Jeschke, J.M., Pagadl, S., Pyšekm, P., van Kleuneno, M., Winterq, M., et al. 2018 Global rise in emerging alien species results from increased accessibility of new source pools. Proc. Natl. Acad. Sci. USA 115, 2264–2273. (doi:10.1073/pnas.1719429115).

[2] van Kleunen, M., Dawson, W., Essl, F., Pergl, J., Winter, M., Weber, E., Kreft, H., Weigelt, P., Kartesz, J., Nishino, M., et al. 2015 Global exchange and accumulation of non-native plants. Nature 525, 100–103. (doi:10.1038/nature14910).

[3] van Kleunen, M., Essl, F., Pergl, J., Brundu, G., Carboni, M., Dullinger, S., Early, R., Gonzalez-Moreno, P., Groom, Q.J., Hulme, P.E., et al. 2018 The changing role of ornamental horticulture in alien plant invasions. Biol. Rev. 93, 1421–1437. (doi:10.1111/brv.12402).

[4] Linders, T.E.W., Schaffner, U., Eschen, R., Abebe, A., Choge, S.K., Nigatu, L., Mbaabu, P.R., Shiferaw, H. & Allan, E. 2019 Direct and indirect effects of invasive species: biodiversity loss is a major mechanism by which an invasive tree affects ecosystem functioning. J. Ecol. 107, 2660–2672. (doi:10.1111/1365-2745.13268).

[5] Vilà, M., Espinar, J.L., Hejda, M., Hulme, P.E., Jarošík, V., Maron, J.L., Pergl, J., Schaffner, U., Sun, Y. & Pyšek, P. 2011 Ecological impacts of invasive alien plants: a meta□analysis of their effects on species, communities and ecosystems. Ecol. Lett. 14, 702–708. (doi:10.1111/j.1461-0248.2011.01628.x).

[6] Pyšek, P., Jarošík, V., Hulme, P.E., Pergl, J., Hejda, M., Schaffner, U. & Vilà, M. 2012 A global assessment of invasive plant impacts on resident species, communities and ecosystems: the interaction of impact measures, invading species’ traits and environment. Glob. Change Biol. 18, 1725–1737. (doi:10.1111/j.1365-2486.2011.02636.x).

[7] Hejda, M., Pyšek, P. & Jarošík, V. 2009 Impact of invasive plants on the species richness, diversity and composition of invaded communities. J. Ecol. 97, 393–403. (doi:10.1111/j.1365-2745.2009.01480.x).

[8] Pyšek, P., Hulme, P.E., Simberloff, D., Bacher, S., Blackburn, T.M., Carlton, J.T., Dawson, W., Essl, F., Foxcroft, L.C., Genovesi, P., et al. 2020 Scientists’ warning on invasive alien species. Biol. Rev. 95, 1511–1534. (doi:10.1111/brv.12627).

[9] Seebens, H., Bacher, S., Blackburn, T.M., Capinha, C., Dawson, W., Dullinger, S., Genovesi, P., Hulme, P.E., van Kleunen, M., Kuhn, I., et al. 2020 Projecting the continental accumulation of alien species through to 2050. Glob. Change Biol. 00, 1–13. (doi:10.1111/gcb.15333).

[10] Essl, F., Lenzner, B., Bacher, S., Bailey, S., Capinha, C., Daehler, C., Dullinger, S., Genovesi, P., Hui, C., Hulme, P.E., et al. 2020 Drivers of future alien species impacts: An expert based assessment. Glob. Change Biol. 26, 4880–4893. (doi:10.1111/gcb.15199).

[11] Bartz, R. & Kowarik, I. 2019 Assessing the environmental impacts of invasive alien plants: a review of assessment approaches. NeoBiota 43, 69–99. (doi:10.3897/neobiota.43.30122).

[12] Liu, Y.J., Oduor, A.M.O., Zhang, Z., Manea, A., Tooth, I.M., Leishman, M.R., Xu, X.L. & van Kleunen, M. 2017 Do invasive alien plants benefit more from global environmental change than native plants? Glob. Change Biol. 23, 3363–3370. (doi:10.1111/gcb.13579).

[13] Dai, A. 2013 Increasing drought under global warming in observations and models. Nat. Clim. Chang. 3, 52–58. (doi:10.1038/NCLIMATE1633).

[14] Spinoni, J., Vogt, J.V., Naumann, G., Barbosa, P. & Dosio, A. 2018 Will drought events become more frequent and severe in Europe? Int. J. Climatol. 38, 1718–1736. (doi:10.1002/joc.5291).

[15] Diffenbaugh, N.S., Swain, D.L. & Touma, D. 2015 Anthropogenic warming has increased drought risk in California. Proc. Natl. Acad. Sci. USA 112, 3931–3936. (doi:10.1073/pnas.1422385112).

[16] Yoon, J.H., Wang, S.Y.S., Gillies, R.R., Kravitz, B., Hipps, L. & Rasch, P.J. 2015 Increasing water cycle extremes in California and in relation to ENSO cycle under global warming. Nat. Commun. 6, 8657. (doi:10.1038/ncomms9657).

[17] Bueno, A., Pritsch, K. & Simon, J. 2020 Responses of native and invasive woody seedlings to combined competition and drought are species-specific. Tree Physiol. 41, 343–357. (doi:10.1093/treephys/tpaa134).

[18] da Silva, E.C., Nogueira, R.J.M.C., da Silva, M.A. & de Albuquerque, M.B. 2011 Drought stress and plant nutrition. Plant stress 5, 32–41.

[19] Manea, A., Sloane, D.R. & Leishman, M.R. 2016 Reductions in native grass biomass associated with drought facilitates the invasion of an exotic grass into a model grassland system. Oecologia 181, 175–183. (doi:10.1007/s00442-016-3553-1).

[20] Werner, C., Zumkier, U., Beyschlag, W. & Máguas, C. 2010 High competitiveness of a resource demanding invasive acacia under low resource supply. Plant Ecol. 206, 83–96. (doi:10.1007/s11258-009-9625-0).

[21] Ahanger, M.A., Morad◻Talab, N., Abd-Allah, E.F., Ahmad, P. & Hajiboland, R. 2016 Plant growth under drought stress: significance of mineral nutrients. Water Stress and Crop Plants: A Sustainable Approach 2, 649–668. (doi:10.1002/9781119054450.ch37).

[22] Haeuser, E., Dawson, W. & van Kleunen, M. 2019 Introduced garden plants are strong competitors of native and alien residents under simulated climate change. J. Ecol. 107, 1328–1342. (doi:10.1111/1365-2745.13101).

[23] Kelso, M.A., Wigginton, R.D. & Grosholz, E.D. 2020 Nutrients mitigate the impacts of extreme drought on plant invasions. Ecology 101, e02980. (doi:10.1002/ecy.2980).

[24] Copeland, S.M., Harrison, S.P., Latimer, A.M., Damschen, E.I., Eskelinen, A.M., Fernandez◻Going, B., Spasojevic, M.J., Anacker, B.L. & Thorne, J.H. 2016 Ecological effects of extreme drought on Californian herbaceous plant communities. Ecol. Monogr. 86, 295–311. (doi:10.1002/ecm.1218).

[25] LaForgia, M.L., Spasojevic, M.J., Case, E.J., Latimer, A.M. & Harrison, S.P. 2018 Seed banks of native forbs, but not exotic grasses, increase during extreme drought. Ecology 99, 896–903. (doi:10.1002/ecy.2160).

[26] Valliere, J.M., Escobedo, E.B., Bucciarelli, G.M., Sharifi, M.R. & Rundel, P.W. 2019 Invasive annuals respond more negatively to drought than native species. New Phytol. 223, 1647–1656. (doi:10.1111/nph.15865).

[27] Pintó-Marijuan, M., Cotado, A., Fleta-Soriano, E. & Munné-Bosch, S. 2017 Drought stress memory in the photosynthetic mechanisms of an invasive CAM species, *Aptenia cordifolia*. Photosynth. Res. 131, 241–253. (doi:10.1007/s11120-016-0313-3).

[28] Eisenhauer, N., Sabais, A.C.W., Schonert, F. & Scheu, S. 2010 Soil arthropods beneficially rather than detrimentally impact plant performance in experimental grassland systems of different diversity. Soil Biol. Biochem. 42, 1418–1424. (doi:10.1016/j.soilbio.2010.05.001).

[29] Bardgett, R.D., Bowman, W.D., Kaufmann, R. & Schmidt, S.K. 2005 A temporal approach to linking aboveground and belowground ecology. Trends Ecol. Evol. 20, 634–641. (doi:10.1016/j.tree.2005.08.005).

[30] Wardle, D.A., Bardgett, R.D., Klironomos, J.N., Setälä, H., van der Putten, W.H. & Wall, D.H. 2004 Ecological linkages between aboveground and belowground biota. Science 304, 1629–1633. (doi:10.1126/science.1094875).

[31] Eisenhauer, N. & Scheu, S. 2008 Earthworms as drivers of the competition between grasses and legumes. Soil Biol. Biochem. 40, 2650–2659. (doi:10.1016/j.soilbio.2008.07.010).

[32] Bardgett, R.D. & Chan, K.F. 1999 Experimental evidence that soil fauna enhance nutrient mineralization and plant nutrient uptake in montane grassland ecosystems. Soil Biol. Biochem. 31, 1007–1014. (doi:10.1016/S0038-0717(99)00014-0).

[33] Barot, S., Ugolini, A. & Brikci, F.B. 2007 Nutrient cycling efficiency explains the long-term effect of ecosystem engineers on primary production. Funct. Ecol. 21, 1–10. (doi:10.1111/j.1365-2435.2006.01225.x).

[34] Marinissen, J.C.Y. & de Ruiter, P.C. 1993 Contribution of earthworms to carbon and nitrogen cycling in agro-ecosystems. Agric. Ecosyst. Environ. 47, 59–74. (doi:10.1016/0167-8809(93)90136-D).

[35] Cole, L., Dromph, K.M., Boaglio, V. & Bardgett, R.D. 2004 Effect of density and species richness of soil mesofauna on nutrient mineralisation and plant growth. Biol. Fertil. Soils 39, 337–343. (doi:10.1007/s00374-003-0702-6).

[36] Smith, V.R. & Steenkamp, M. 1992 Soil macrofauna and nitrogen on a sub-Antarctic island. Oecologia 92, 201–206. (doi:10.1007/BF00317365).

[37] Xu, G.L., Mo, J.G., Zhou, G.Y. & Peng, S.L. 2003 Relationship of soil fauna and N cycling and its response to N deposition. Acta Ecologica Sinica 23, 2453–2463. (doi:10.3321/j.issn:1000-0933.2003.11.030).

[38] Liu, Y.J., Liu, M., Xu, X.L., Tian, Y.Q., Zhang, Z. & van Kleunen, M. 2018 The effects of changes in water and nitrogen availability on alien plant invasion into a stand of a native grassland species. Oecologia 188, 441–450. (doi:10.1007/s00442-018-4216-1).

[39] Bonkowski, M., Villenave, C. & Griffiths, B. 2009 Rhizosphere fauna: the functional and structural diversity of intimate interactions of soil fauna with plant roots. Plant Soil 321, 213–233. (doi:10.1007/s11104-009-0013-2).

[40] Korell, L., Schädler, M., Brandl, R., Schreiter, S. & Auge, H. 2019 Release from above- and belowground insect herbivory mediates invasion dynamics and impact of an exotic plant. Plants 8, 544. (doi:10.3390/plants8120544).

[41] Keane, R.M. & Crawley, M.J. 2002 Exotic plant invasions and the enemy release hypothesis. Trends Ecol. Evol. 17, 164–170. (doi:10.1016/S0169-5347(02)02499-0).

[42] Liu, H. & Stiling, P. 2006 Testing the enemy release hypothesis: a review and meta-analysis. Biol. Invasions 8, 1535–1545. (doi:10.1007/s10530-005-5845-y).

[43] Mitchell, C.E. & Power, A.G. 2003 Release of invasive plants from fungal and viral pathogens. Nature 421, 625–627. (doi:10.1038/nature01317).

[44] Vilà, M., Maron, J.L. & Marco, L. 2005 Evidence for the enemy release hypothesis in *Hypericum perforatum*. Oecologia 142, 474–479. (doi:10.1007/s00442-004-1731-z).

[45] Ockendon, N., Baker, D.J., Carr, J.A., White, E.C., Almond, R.E.A., Amano, T., Bertram, E., Bradbury, R.B., Bradley, C., Butchart, S.H., et al. 2014 Mechanisms underpinning climatic impacts on natural populations: altered species interactions are more important than direct effects. Glob. Change Biol. 20, 2221–2229. (doi:10.1111/gcb.12559).

[46] Franco, A.L.C., Gherardi, L.A., de Tomasel, C.M., Andriuzzi, W.S., Ankrom, K.E., Bach, E.M., Guan, P., Sala, O.E. & Wall, D.H. 2020 Root herbivory controls the effects of water availability on the partitioning between above◻and below◻ground grass biomass. Funct. Ecol. 34, 2403–2410. (doi:10.1111/1365-2435.13661).

[47] Guyer, A., Hibbard, B.E., Holzkämper, A., Erb, M. & Robert, C.A.M. 2018 Influence of drought on plant performance through changes in belowground tritrophic interactions. Ecol. Evol. 8, 6756–6765. (doi:10.1002/ece3.4183).

[48] Erb, M., Köllner, T.G., Degenhardt, J., Zwahlen, C., Hibbard, B.E. & Turlings, T.C.J. 2011 The role of abscisic acid and water stress in root herbivore-induced leaf resistance. New Phytol. 189, 308–320. (doi:10.1111/j.1469-8137.2010.03450.x).

[49] Wilschut, R.A. & van Kleunen, M. 2021 Drought alters plant◻soil feedback effects on biomass allocation but not on plant performance. Plant Soil 462, 285–296. (doi:10.1007/s11104-021-04861-9).

[50] Aupic-Samain, A., Baldy, V., Delcourt, N., Krogh, P.H., Gauquelin, T., Fernandez, C. & Santonja, M. 2020 Water availability rather than temperature control soil fauna community structure and prey◻predator interactions. Funct. Ecol. (doi:10.1111/1365-2435.13745).

[51] Eisenhauer, N., Cesarz, S., Koller, R., Worm, K. & Reich, P.B. 2012 Global change belowground: impacts of elevated CO_2_, nitrogen, and summer drought on soil food webs and biodiversity. Glob. Change Biol. 18, 435–447. (doi:10.1111/j.1365-2486.2011.02555.x).

[52] Makkonen, M., Berg, M.P., Van Hal, J.R., Callaghan, T.V., Press, M.C. & Aerts, R. 2011 Traits explain the responses of a sub-arctic collembola community to climate manipulation. Soil Biol. Biochem. 43, 377–384. (doi:10.1016/j.soilbio.2010.11.004).

[53] Wilschut, R.A. & Geisen, S. 2021 Nematodes as drivers of plant performance in natural systems. Trends Plant Sci. 26, 237–247. (doi:10.1016/j.tplants.2020.10.006).

[54] Yan, X.L., Wang, Z.H. & Ma, J.S. 2019 The checklist of the naturalized plants in china. Shanghai, China, Shanghai Scientific and Technical Publishers.

[55] Parepa, M., Fischer, M. & Bossdorf, O. 2013 Environmental variability promotes plant invasion. Nat. Commun. 4, 1604. (doi:10.1038/ncomms2632).

[56] Pinheiro, J., Bates, D., DebRoy, S., Sarkar, D. & R Core Team. 2020 nlme: linear and nonlinear mixed effects models. (R package version 3.1-151.

[57] R Core Team. 2020 R: A language and environment for statistical computing. In R Foundation for Statistical Computing (Vienna, Austria.

[58] Zuur, A.F., Ieno, E.N., Walker, N.J., Saveliev, A.A. & Smith, G.M. 2009 Mixed effects models and extensions in ecology with R. New York, USA, Springer.

[59] Zlatev, Z. & Lidon, F.C. 2012 An overview on drought induced changes in plant growth, water relations and photosynthesis. Emir. J. Food Agric. 24, 57–72. (doi:10.1111/j.1439-0426.2005.00664.x).

[60] Beierkuhnlein, C., Thiel, D., Jentsch, A., Willner, E. & Kreyling, J. 2011 Ecotypes of European grass species respond differently to warming and extreme drought. J. Ecol. 99, 703–713. (doi:10.1111/j.1365-2745.2011.01809.x).

[61] Gupta, A., Rico-Medina, A. & Caño-Delgado, A.I. 2020 The physiology of plant responses to drought. Science 368, 266–269. (doi:10.1126/science.aaz7614).

[62] Richards, C.L., Bossdorf, O., Muth, N.Z., Gurevitch, J. & Pigliucci, M. 2006 Jack of all trades, master of some? On the role of phenotypic plasticity in plant invasions. Ecol. Lett. 9, 981–993. (doi:10.1111/j.1461-0248.2006.00950.x).

[63] Lussenhop, J. & Bassirirad, H. 2005 Collembola effects on plant mass and nitrogen acquisition by ash seedlings (*Fraxinus pennsylvanica*). Soil Biol. Biochem. 37, 645–650. (doi:10.1016/j.soilbio.2004.08.021).

[64] Mehring, A.S. & Levin, L.A. 2015 Potential roles of soil fauna in improving the efficiency of rain gardens used as natural stormwater treatment systems. J. Appl. Ecol. 52, 1445–1454. (doi:10.1111/1365-2664.12525).

[65] Partsch, S., Milcu, A. & Scheu, S. 2006 Decomposers (lumbricidae, collembola) affect plant performance in model grasslands of different diversity. Ecology 87, 2548–2558. (doi:10.1890/0012-9658(2006)87[2548:DLCAPP]2.0.CO;2).

[66] Blossey, B. & Notzold, R. 1995 Evolution of increased competitive ability in invasive nonindigenous plants: a hypothesis. J. Ecol. 83, 887–889. (doi:10.2307/2261425).

[67] Chun, Y.J., van Kleunen, M. & Dawson, W. 2010 The role of enemy release, tolerance and resistance in plant invasions: linking damage to performance. Ecol. Lett. 13, 937–946. (doi:10.1111/j.1461-0248.2010.01498.x).

[68] Ramula, S., Paige, K.N., Lennartsson, T. & Tuomi, J. 2019 Overcompensation: a 30-year perspective. Ecology 100, e02667. (doi:10.1002/ecy.2667).

[69] Garcia, L.C. & Eubanks, M.D. 2019 Overcompensation for insect herbivory: a review and meta-analysis of the evidence. Ecology 100, e02585. (doi:10.1002/ecy.2585).

[70] Gange, A. 2000 Arbuscular mycorrhizal fungi, collembola and plant growth. Trends Ecol. Evol. 15, 369–372. (doi:10.1016/S0169-5347(00)01940-6).

[71] de Sassi, C. & Tylianakis, J.M. 2012 Climate change disproportionately increases herbivore over plant or parasitoid biomass. PLoS One 7, e40557. (doi:10.1371/journal.pone.0040557).

[72] Meza-Lopez, M.M. & Siemann, E. 2020 Warming alone increased exotic snail reproduction and together with eutrophication influenced snail growth in native wetlands but did not impact plants. Sci. Total Environ. 704, 135271. (doi:10.1016/j.scitotenv.2019.135271).

[73] Nooten, S.S. & Hughes, L. 2014 Potential impacts of climate change on patterns of insect herbivory on understorey plant species: a transplant experiment. Austral Ecol. 39, 668–676. (doi:10.1111/aec.12129).

[74] Classen, A.T., Sundqvist, M.K., Henning, J.A., Newman, G.S., Moore, J.A.M., Cregger, M.A., Moorhead, L.C. & Patterson, C.M. 2015 Direct and indirect effects of climate change on soil microbial and soil microbial-plant interactions: What lies ahead? Ecosphere 6, 130. (doi:10.1890/es15-00217.1).

[75] Kardol, P., Cregger, M.A., Campany, C.E. & Classen, A.T. 2010 Soil ecosystem functioning under climate change: plant species and community effects. Ecology 91, 767–781. (doi:10.1890/09-0135.1).

[76] Lindberg, N. 2003 Soil fauna and global change. Niklas Lindberg, Uppsala, Sweden.

[77] Levine, M.T. & Paige, K.N. 2004 Direct and indirect effects of drought on compensation following herbivory in Scarlet gilia. Ecology 85, 3185–3191. (doi:10.1890/03-0748).

